# Differential induction of 14-3-3 paralogs expression during early and late adipogenesis

**DOI:** 10.64898/2026.08.03.742103

**Authors:** Samanta del Veliz, Sergio Müller, Jonathan N. Aguilera, Aldana Daniela Gojanovich, Marina Uhart, Gareth Lim, Diego M Bustos

## Abstract

Obesity is a major public health challenge of the 21st century, particularly in low- and middle-income populations. Adipogenesis plays a central role in the development of obesity and associated metabolic disorders, as it determines adipocyte number, size, and function. The 14-3-3 protein family comprises seven paralogs in mammals that regulate multiple cellular processes, yet their specific roles during adipogenesis remain poorly understood. In this study, we characterized the expression profiles of 14-3-3 paralogs during the early and late stages of adipogenic differentiation using quantitative PCR under standard adipogenic differentiation medium and modified drug-supplemented conditions. We found that the expression of specific paralogs is strongly influenced by the composition of the differentiation medium. The absence of insulin led to an early increase in *Ywhaz*, which could not be maintained during late adipogenesis and was associated with impaired adipogenic differentiation. In contrast, stimulation with incretins in combination with insulin induced late expression of *Ywhag* and *Ywhab* paralogs and promoted the formation of a greater number of smaller lipid droplets. These findings indicate that individual 14-3-3 paralogs exert distinct and context-dependent effects on adipogenesis, highlighting their potential roles as modulators of adipocyte differentiation and metabolic function.

## Introduction

The increase in adipocyte number and size occurs through adipocyte differentiation (adipogenesis), a process crucial for the development of obesity (Rial et al., 2024). Obesity is characterized by adipose tissue expansion driven by adipocyte hypertrophy and hyperplasia, processes dependent on adipogenesis (Westbury et al., 2023).

Adipogenesis contributes to obesity and associated metabolic disorders. This process is sensitive to multiple factors that influence adipocyte size (hypertrophy), number (hyperplasia), adipocyte function, and paracrine effects, including the local action of cytokines and adipokines secreted by adipocytes and immune cells, which regulate differentiation, proliferation, and inflammation within adipose tissue.

Adipocyte differentiation is initiated by extracellular signals received by mesenchymal or stromal stem cells. In vitro induction of adipogenesis can be facilitated using drugs that modulate specific signaling pathways. For example, rosiglitazone acts as an agonist of the nuclear receptor PPAR-γ, which is critical for the formation of mature adipocytes (Fayyad et al., 2019). Insulin increases the expression of PPAR-γ and other lipogenesis-related genes (Ambele et al., 2020), while dexamethasone regulates PPAR-γ and C/EBPδ, a factor involved in the early expression of pro-adipogenic genes (Asada et al., 2011; Akavia et al., 2006; Lee et al., 2014). 3-Isobutyl-1-methylxanthine (IBMX) increases intracellular cAMP and activates the PKA pathway, promoting the expression of adipogenic genes. A similar agent is glucagon-like peptide 1 (GLP-1), which also activates the PKA pathway and elevates intracellular cAMP. However, GLP-1 receptor activation triggers signaling pathways that interfere with differentiation, suggesting that cAMP elevation alone is not sufficient to recapitulate IBMX- induced adipogenic effects (Lee et al., 2015; Liu et al., 2020).

It is well established that 14-3-3 proteins regulate adipogenesis through multiple mechanisms, including direct interaction with Lipin-1 and modulation of chromatin remodeling. In adipocytes, Lipin-1 was shown to bind 14-3-3, a binding that governs its subcellular localization: overexpression of 14-3-3 promotes Lipin-1 retention in the cytoplasm, a regulation that depends on several phosphorylated serine/threonine residues within Lipin-1’s serine-rich domain (Péterfy et al., 2010).

Furthermore, recent proteomic and chromatin-accessibility studies demonstrated that the nuclear interactome of 14-3-3ζ is enriched in chromatin-modifying enzymes (e.g., DNMT1, HDAC1), and depletion of 14-3-3ζ significantly alters chromatin accessibility across hundreds of genomic regions corresponding to adipogenic genes (Rial et al, 2024).

Importantly, 14-3-3ζ knockdown impairs the induction of master adipogenic transcription factors — including C/EBP-δ, PPARγ and C/EBP-α — and blocks adipocyte differentiation both in vitro and in vivo (Lim et al., 2015).

Altogether, these data support a model in which 14-3-3, acts upstream of the adipogenic transcriptional program and is critical for the proper expression of key regulators of adipocyte maturation. The expression of 14-3-3γ and β paralogs increases during differentiation of 3T3-L1 preadipocytes, and similar results have been observed for 14-3-3ζ in correlation with intracellular lipid accumulation (Gojanovich et al., 2016; Lim S Johnson, 2016). Certain differentiation-promoting drugs activate signaling pathways in which 14-3-3 proteins play a role. For example, 14-3-3 proteins are essential in every phase of insulin signaling, from the initiation of signal transduction to glucose uptake and subsequent transcriptional responses (Pennington et al., 2018). However, there is little information on how these drugs, used in vitro or as in vivo therapeutics, modulate 14-3-3 expression. In this study, we characterized the expression of 14-3-3γ, β, and ζ during early and late adipogenesis in response to different adipogenic drugs, correlating these findings with the degree of adipocyte differentiation. These results provide insight into the differential regulation of 14-3-3 paralogs during adipocyte differentiation and their association with drug-specific adipogenic signaling pathways.

## Material and Methods

### 1. Cell culture, adipogenic differentiation and treatments

The NIH 3T3-L1 mouse preadipocyte cell line (CL-173, ATCC) was routinely grown in 10 cm diameter culture dishes in complete Dulbecco’s modified Eagle medium - 12800-017, Gibco Laboratories, Thermo Fisher Scientific (DMEM), with high glucose (4.5 g/L), 0.5 mM sodium pyruvate and glutamine. Complete DMEM was prepared through supplementation with 10% Fetal Bovine Serum (FBS Internegocios), 100 U/mL penicillin, and 100 *μ*g/mL streptomycin (Thermo Fisher Scientific). Cells were maintained in a 5% CO_2_ atmosphere in an incubator at 37 °C (Gojanovich et al. 2016b).

Cells were seeded on a six-well plate at a density of 3 x 10^3^ cells per cm^2^ and grown in DMEM to 80% confluence. To induce adipogenic differentiation, media was removed, and the following different combinations of differentiation media were added in a specific timeline:

Then, we carried out a sample collection for RNA to study the expression of *Ywha* genes, which encode for the different 14-3-3 paralogs, during early adipogenesis (day 3). To evaluate late adipogenesis, samples were obtained for qPCR on day 7 of differentiation, in which the cells already showed an accumulation of lipid droplets, which were visualized by *Oil Red O* staining (Gojanovich et al. 2018).

The cell lines stably expressing shRNAs targeting 14-3-3γ (YWHAG) or 14-3-3β (YWHAB) were previously established, validated, and characterized in our laboratory, as described in Frontini-López et al. 2021 and Rivera et al. 2026.

### 2. RNA extraction, cDNA production, and qPCR

Purification of total RNA from UT 3T3-L1 and differentiated cells was done following the manufacturer’s instructions. Cells were grown in a 6-well plate and when they reached day 3 or 7 post-induction, 1 mL of RNA extraction reagent Bio-Zol (RA02, Productos Bio- Lógicos, Argentina) was added to each sample. Tubes were centrifuged at 12,000 x *g* for 10 min at 4 °C and then the upper phase, containing the total RNA, was transferred to a new tube. RNA was precipitated by adding 500 *μ*L of isopropanol and incubated for 1 hour at -20 °C. Subsequently, the tubes were centrifuged at 12,000 x *g* for 10 min at 4 °C to obtain an RNA pellet. The pellet was dissolved in 25 *μ*L of RNase-free water. The amount of RNA obtained was quantified by spectrophotometry and its integrity was observed by 2% agarose gel. The RNA was stored at -80 °C until use. For cDNA synthesis by reverse transcription, cDNA synthesis was performed from 1 *μ*g of RNA per sample. 1 U of DNase I (EN0521, Thermo Fisher Scientific, USA) was added. Reverse transcription was performed by adding 2 *μ*L of 10 *μ*M random hexamers, 2 *μ*L of dNTPs and nuclease-free water to complete a final volume of 15 *μ*L. The samples were heated at 70 °C for 5 min with 1 *μ*L (200 U) of the M-MLV Transcriptase enzyme (EA13, Productos Bio-Lógicos, Argentina) and 4 *μ*L of 5X reaction buffer provided with the enzyme. A first incubation was done at 25 °C for 10 min and then at 37 °C for one hour. Samples were diluted to a final volume of 50 *μ*L using nuclease-free water and stored at -20 °C until use. For quantitative real-time PCR, primers were chosen by amplicon size (between 100 and 300 bp), and care was taken that their sequences hybridized in exons on both sides of an intron (intron spanning), ensuring that the products were cDNA-specific. For β-actin (ACTB), we used primers previously published by our laboratory (Gojanovich et al., 2016b). The sequences of the primers used in this work are described in Table 1 and were synthesized by the company Macrogen (South Korea). A Master mix reaction tube was prepared using Maxima SYBR Green qPCR Master Mix (2X) (K0251, Thermo Fisher Scientific, USA), 0.3 *μ*M Forward primer (F), 0.3 *μ* M Reverse primer (R), and 10 nM ROX solution (R1371, Thermo Fisher Scientific, USA). The QuantStudio^TM^ 6 Flex Real-Time PCR System thermocycler (4485691, Thermo Fisher Scientific, USA) was used. The relative expression of each gene was analyzed based on the method of Livak and Schmittgen (2001). The fold-change in the expression of the genes of interest for each condition was represented by the calculation of 2^(-ΔΔCt)^.

**Table 1:** Timeline and medium composition for differentiation of 3T3-L1 cells.

| Condition / Medium | Day 0–3 | Day 3–5<br>(Maintenance) | Day 5–7<br>(Final) | Key Additives |
| --- | --- | --- | --- | --- |
| A. ADM (Adipogenic Differentiation Medium) | Complete DMEM + 0.5 mM IBMX + 2.5 µM Dexamethasone + 2 µM Rosiglitazone + 10 µg/mL Insulin | Complete DMEM + 10 µg/mL Insulin | Complete DMEM | IBMX, Dexamethasone, Rosiglitazone, Insulin |
| B. ADM-IBMX+GLP-1 | Complete DMEM + 1 µg/mL Incretin + 2.5 µM Dexamethasone + 2 µM Rosiglitazone + 10 µg/mL Insulin | Complete DMEM + 10 µg/mL Insulin | Complete DMEM | Incretin replaces IBMX; Dexamethasone, Rosiglitazone, Insulin |
| **C. ADM-insulin+GLP-1** | Complete DMEM + 0.5 mM IBMX + 2.5 μM Dexamethasone + 2 μM Rosiglitazone + 1 μg/mL Incretin | Complete DMEM + 1 μg/mL GLP-1 | Complete DMEM | Incretin replaces Insulin; IBMX, Dexamethasone, Rosiglitazone |
| **D. ADM-insulin** | Complete DMEM + 0.5 mM IBMX + 2.5 μM Dexamethasone + 2 μM Rosiglitazone | Complete DMEM + 10% FBS | Complete DMEM + 10% FBS | No Insulin; IBMX, Dexamethasone, Rosiglitazone |
| **E. Insulin Medium** | Complete DMEM + 10 μg/mL Insulin | Complete DMEM + 10 μg/mL Insulin | Complete DMEM | Insulin only |
| **F. Untreated Cells (UT)** | Complete DMEM + 2.5% FBS | Complete DMEM + 2.5% FBS | Complete DMEM + 2.5% FBS |  |

### 3. Red Oil O staining

3T3-L1 cells differentiated into adipocytes and their corresponding negative control samples were stained with *Oil Red O* (Biopack, Buenos Aires, Argentina) (Gojanovich et al. 2018.) This dye stains specifically the neutral lipids (triglycerides and cholesterol oleate) contained in the lipid droplets (LDs). A stock solution 0.35 % w/v in isopropanol, filtered first through 0.45 *μ*m nitrocellulose membrane and then through 0.22 *μ*m was prepared and stored at room temperature for at least 24 hours. After 7 days of differentiation, the treated and untreated cells attached to circular 12 mm diameter coverslips were washed three times with PBS containing 1 mM CaCl_2_ and 1 mM MgCl_2_. The cells were fixed with 4% w/v PFA (paraformaldehyde; Sigma-Aldrich, USA) for 20 min at room temperature and then washed 3 times with PBS. The stock solution was diluted in Milli-Q water (in a ratio of 3 parts of the stock solution: 2 parts of Milli-Q water) and filtered again through a 0.22 *μ*m nitrocellulose membrane, obtaining a working solution. The latter was added (approximately 300 *μ*L per well of 24 wells plate) and incubated for 2 hours at room temperature with gentle shaking and protected from light. The samples were washed with Milli-Q water and mounted on slides with 8 *μ*L Mowiol 4-88 (Sigma-Aldrich) for optical microscopy analysis.

### 4. SDS-PAGE and Western blots

Proteins were separated by SDS–PAGE using 12% polyacrylamide gels and transferred onto PVDF membranes using Towbin transfer buffer (25 mM Tris, 192 mM glycine, 0.1% SDS, and 20% v/v methanol) at 10 V overnight at 4 °C. Membranes were blocked with 5% milk in PBS for 1 h and washed three times with 1× PBS for 5 min each. Primary antibody incubation was performed for 60 min in PBS containing 3% BSA using the following antibodies: anti-14-3-3 β (C-20, sc-628; 1:100 Santa Cruz), anti-14-3-3 γ (Martin *et al*.; 1:500), and anti-β-tubulin (Sigma-Aldrich; 1:1000). After three 5-min washes with 0.01% PBS-Tween, membranes were incubated with secondary antibodies diluted in 0.01% PBS-Tween containing 5% milk: bovine anti-rabbit IgG-HRP (sc-2370; 1:3000 Santa Cruz) for 14-3-3 γ and β, and horse anti-mouse IgG-HRP (H+L, PI-2000-1; 1:2000) for β-tubulin. Following three additional 5-min washes with 0.01% PBS-Tween and a final rinse with 1× PBS, signal detection was performed using the Clarity™ Western ECL Substrate (Bio-Rad). Band intensities were quantified using Image Studio software (LI-COR).

### 5. Data Presentation and Statistical Analysis

Data are presented as mean ± standard deviation of the mean (SD) using Graphad Prism. Statistical significance was calculated by one-way ANOVA followed by Dunnett’s, Tukey or Bonferroni test as appropriate. The data were considered statistically significant when *p* < 0.05.

## Results

### 1. Relative expression of *Ywha* paralogs during early and late adipogenesis

We analyzed mRNA expression of the seven murine 14-3-3 paralogs during early (day 3 after adipogenesis induction) and late (day 7) adipogenesis. mRNA was isolated from untreated 3T3-L1 control cells (UT) and cells induced to undergo adipogenic differentiation by incubation with Adipogenic Differentiation Media (ADM, n = 8). Relative expression of the seven *Ywha* paralogs on day 3 of adipogenic differentiation revealed a significant (≥2.0-fold) upregulation of three paralogs – *Ywha z*, *g*, and *b* – in 3T3-L1 cells (Fig. 1A). The higher expression levels of these paralogs were increased at day 7 of differentiation (≥3.0-fold, Fig. 1B).

**Figure 1.**
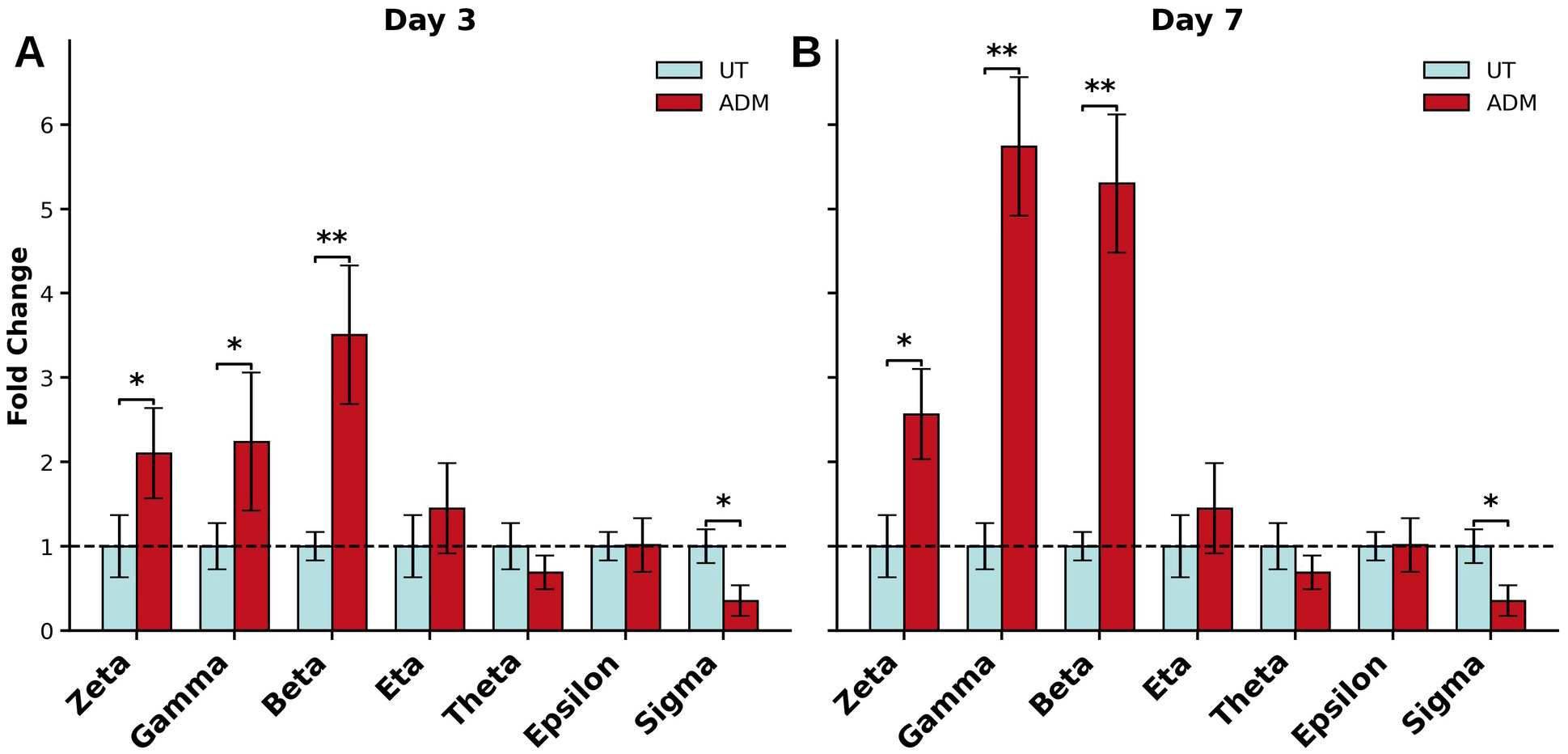
Relative expression levels (fold change) of the seven 14-3-3 paralog groups (Zeta, Gamma, Beta, Eta, Theta, Epsilon, and Sigma) under untreated conditions (UT, light blue) and ADM treatment (ADM, red). (A) Day 3 and (B) Day 7 post-treatment. Expression values were normalized to the UT control, represented by the dashed horizontal line at fold change = 1. Bars represent mean ± SD. Statistical analysis was performed using one-way ANOVA followed by post hoc multiple-comparison testing. Significant differences between UT and ADM conditions are indicated by asterisks (* p < 0.05, ** p < 0.01). ADM treatment induced marked upregulation of Zeta, Gamma, and Beta paralogs, particularly at Day 7, whereas Sigma expression was significantly reduced under ADM conditions.

Among all *Ywha* paralogs, *g* and *b* showed the strongest induction (Fig 1A and 1B) exhibiting up to ≥ 5-fold increase.

### 2. Relative expression of *Ywha z*, g and *b* on different adipogenesis induction media

We next examined the effects of different compounds on *Ywhaz, Ywhag*, and *Ywhab* mRNA levels in 3T3-L1 cells undergoing adipogenesis.

The figure 2 shows qPCR-based fold changes in mRNA expression (normalized to actin) for the *Ywhaz, Ywhag*, and *Ywhab* paralogs under different treatment conditions at day 3 and day 7. In Figure 2A, *Ywhaz* mRNA levels increased following the induction of adipogenesis under all conditions. No significant differences in *Ywhaz* expression were observed between conditions at either day 3 or day 7. However, under conditions in which insulin was absent, *Ywhaz* expression decreased from day 3 to day 7, whereas this decline was not observed in the presence of insulin. Because insulin was the only variable differing between conditions, these results suggest that it contributes to maintaining *Ywhaz* expression during adipogenic differentiation.

**Figure 2.**
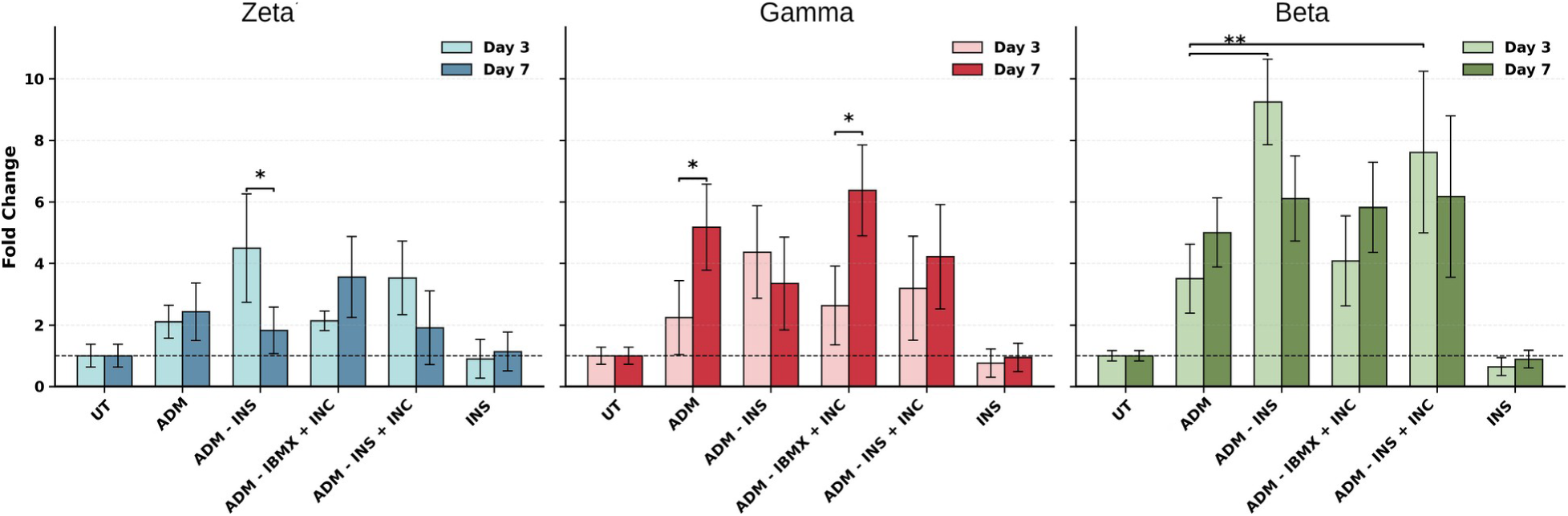
Relative expression levels (fold change) of the Zeta, Gamma, and Beta 14-3-3 paralogs under different treatment conditions at Day 3 and Day 7. Expression levels were normalized to the untreated control (UT), indicated by the dashed horizontal line at fold change = 1. Treatments included MDA, ADM − INS, ADM − IBMX + INC, ADM − INS + INC, and INS. Bars represent mean ± SD. Statistical analysis was performed using one-way ANOVA followed by post hoc multiple-comparison testing. Significant differences between Day 3 and Day 7 within the same treatment are indicated by asterisks (* p < 0.05, ** p < 0.01). Gamma and Beta paralogs showed the strongest induction under ADM-related treatments, whereas Zeta displayed a more moderate and variable response depending on treatment conditions.

In contrast, *Ywhag* displayed a distinct expression pattern (Fig. 2B). Its expression was induced during adipogenic differentiation with ADM. Under insulin-deprived conditions, *Ywhag* expression was not reduced at day 7, unlike *Ywhaz*. The expression pattern observed with ADM was also observed in ADM lacking IBMX but supplemented with incretin, indicating that cAMP-mediated signaling is required for both the induction and maintenance of *Ywhag* expression, independent of the upstream mechanisms driving its induction.

For *Ywhab*, the expression pattern was similar to that observed for *Ywhaz* (Fig. 2A and 2C), although with greater differences between conditions. In the absence of insulin, its expression was significantly higher at day 3, with levels declining by day 7 to those observed under standard ADM conditions. Expression levels under conditions supplemented with IBMX or incretins were comparable.

Treatment with complete medium supplemented only with insulin produced expression levels of *Ywhaz*, *g* and *b* similar to those in untreated cells, suggesting that insulin alone is not sufficient to induce the up-regulation of their mRNAs.

To confirm incretins activity in 3T3-L1 cells, we used a cAMP sensor plasmid, *pcDNA3- AKAP79-(Ci/Ce)-Epac2-camps* (Tener *et al*., 2022). Cells were transfected and visualized under a confocal microscope. Upon addition of incretin (1 µM), green fluorescence intensity markedly decreased compared to untreated controls, consistent with an increase in intracellular cAMP levels (Fig. 3).

**Figure 3.**
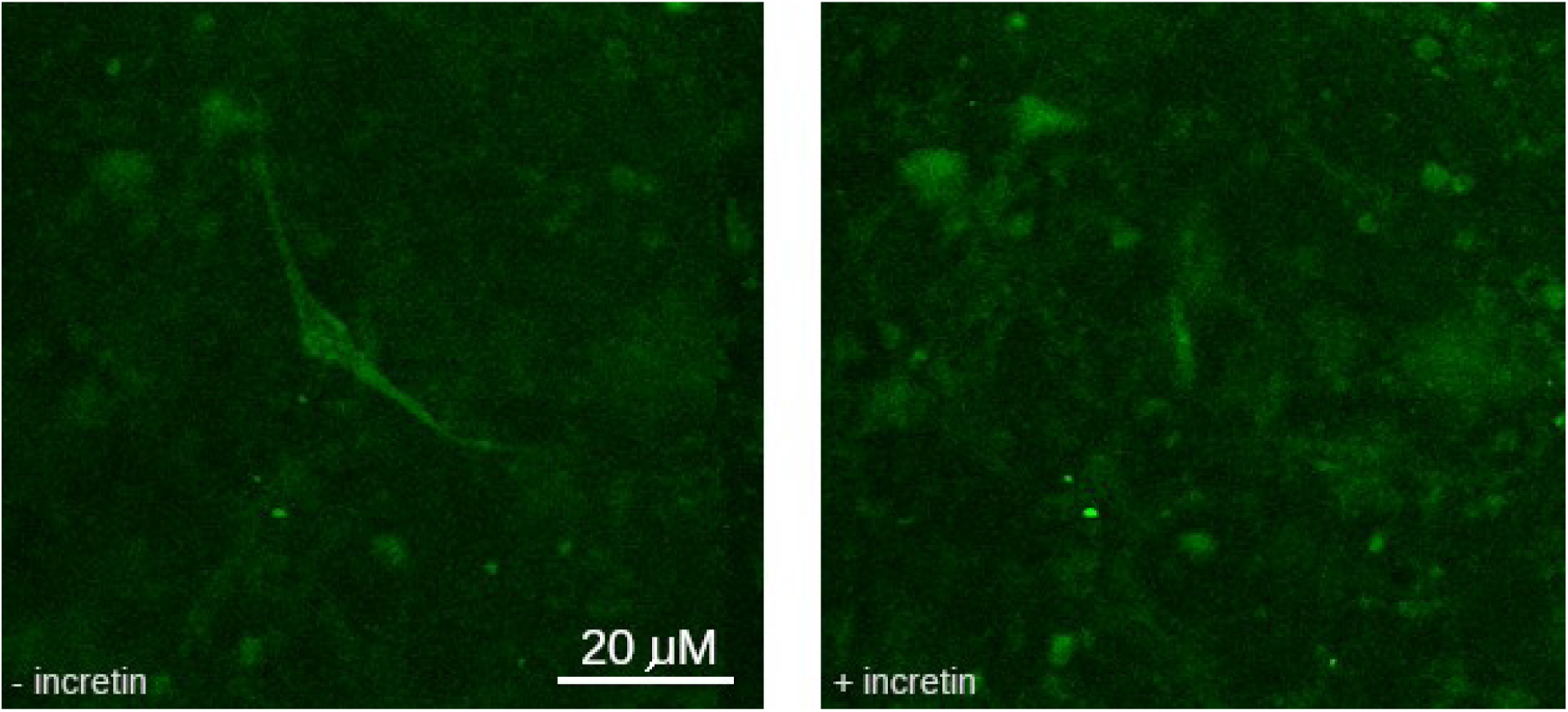
Representative fluorescence images of cells transiently transfected with the cAMP FRET biosensor pcDNA3-AKAP79-(Ci/Ce)-Epac2-camps (Tener et al., 2022). Cells were differentiated in the absence (− incretin) or presence (+ incretin) of incretins. Representative images show the fluorescence distribution of the biosensor under both differentiation conditions. Scale bar: XX μm.

### 3. Insulin influences differently the protein levels of 14-3-3γ and 14-3-3β

Considering the opposite effects of insulin on *Ywhag* and *Ywhab* at the mRNA level, it was of interest to evaluate its influence at the protein level under conditions in which their mRNA levels remained unchanged. To this end, cells were incubated in complete DMEM supplemented with different concentrations of insulin for a short period (6 hours), allowing us to determine whether protein levels also remained stable.

As shown in Figure 4, 14-3-3γ expression remained relatively constant across untreated (UT), 5 µg/ml insulin (lane 1), and 10 µg/ml insulin (lane 2), with a moderate increase observed at 15 µg/ml insulin (lane 3), compatible with the mRNA data. In contrast, 14-3-3β levels decreased markedly and progressively from the UT condition to 15 µg/ml insulin, where it was nearly undetectable under our experimental conditions.

**Figure 4.**
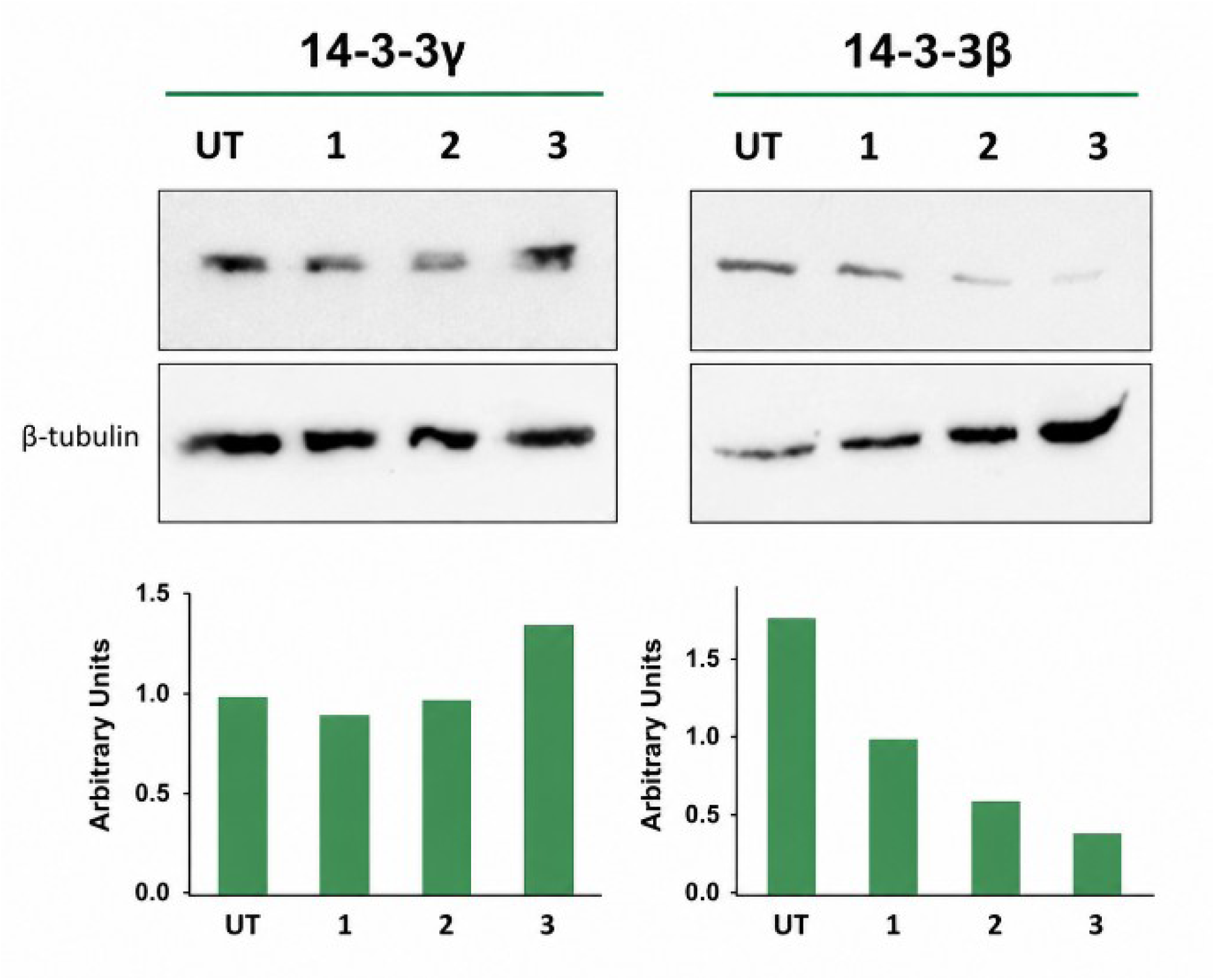
Western blot analysis of 14-3-3γ and 14-3-3β protein levels in non-treated cells (NT), and cells treated with insulin 5 µg/ml (1), 10 µg/ml (2), and 15 µg/ml (3) during 6 hours. Upper panels show representative immunoblots for 14-3-3γ (left) and 14-3-3β (right), with β-tubulin used as a loading control. The corresponding bar graphs below display the densitometric quantification of each band normalized to β-tubulin and expressed in arbitrary units.

These results indicate that insulin differential regulates 14-3-3 paralogs at the protein level, in agreement with their distinct transcriptional responses observed during adipogenesis.

### 4. Drug combinations in differentiation media affect the accumulation of lipid droplets in 3T3- L1 cells

To correlate *Ywha* paralog expression and adipogenic differentiation of 3T3-L1 cells, we stained the cells with *Oil Red O* on day 7 of differentiation, (lipid accumulation at day 3 is not enough to be detected by *Oil Red O*). The degree of differentiation was confirmed by quantifying the red pixels of the images obtained in bright field microscopy, using an *in-house* software developed by us (Masone *et al*. 2017). Figure 5 shows microscopy images of all differentiation conditions, compared to untreated 3T3-L1 cells. Cells treated with all tested drug combinations showed significant differences in accumulation of lipid droplets (LDs) compared to untreated control cells.

**Figure 5.**
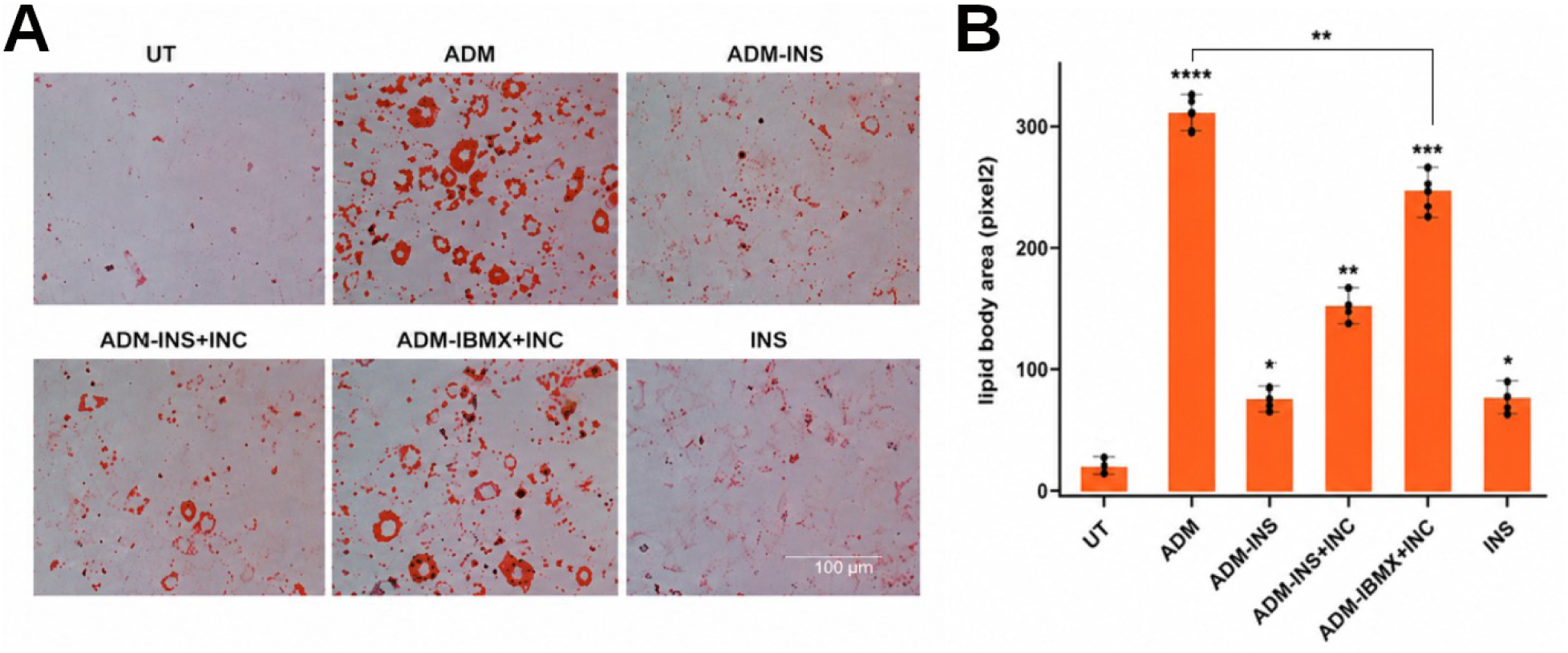
Bright-field micrographs of 3T3-L1 cells stained with Oil Red O (A) and their quantification (B). Cells were grown in DMEM + 10% FBS, ADM, ADM without IBMX plus incretins, ADM without insulin plus incretins, ADM without insulin and IBMX, DMEM with insulin. The statistical test used was the Two-Sample T-test (Welch’s T-test). Significant differences were marked as follows: against UT * p < 0.05, ** p < 0.01, *** p < 0.001, **** p < 0.0001.

Oil Red O staining revealed that ADM-treated cells (positive control, Fig. 5B) displayed the most pronounced lipid accumulation, showing the largest and most abundant LDs compared to untreated cells. In contrast, insulin alone and ADM–insulin conditions resulted in the lowest LD accumulation, both significantly reduced relative to the ADM positive control.

Interestingly, ADM–insulin+incretin treatment led to a marked increase in lipid accumulation -though not as high as ADM- suggesting that incretins can partially rescue differentiation in the absence of insulin. Finally, replacing IBMX with incretin (ADM– IBMX+incretin) also promoted LD formation; however, lipid droplets were smaller and more numerous. Quantitative analysis indicated that differentiation under this condition was slightly lower than under ADM, but still significantly higher than in untreated cells (Fig. 5B).

### 5. Expression of 14-3-3γ, β and ζ paralogs and the LDs accumulation

The figure 6 summarizes the relationship between the level of expression of the three 14-3- 3 paralogs studied here and the stage of differentiation of the 3T3-L1 cells. This scheme shows the differentiation levels generated in 3T3-L1 cells using the assayed differentiation media. Also, it compares how 14-3-3*γ*, *β* and *ζ* paralogs significantly increase during the early and late stages of differentiation compared to undifferentiated cells.

**Figure 6.**
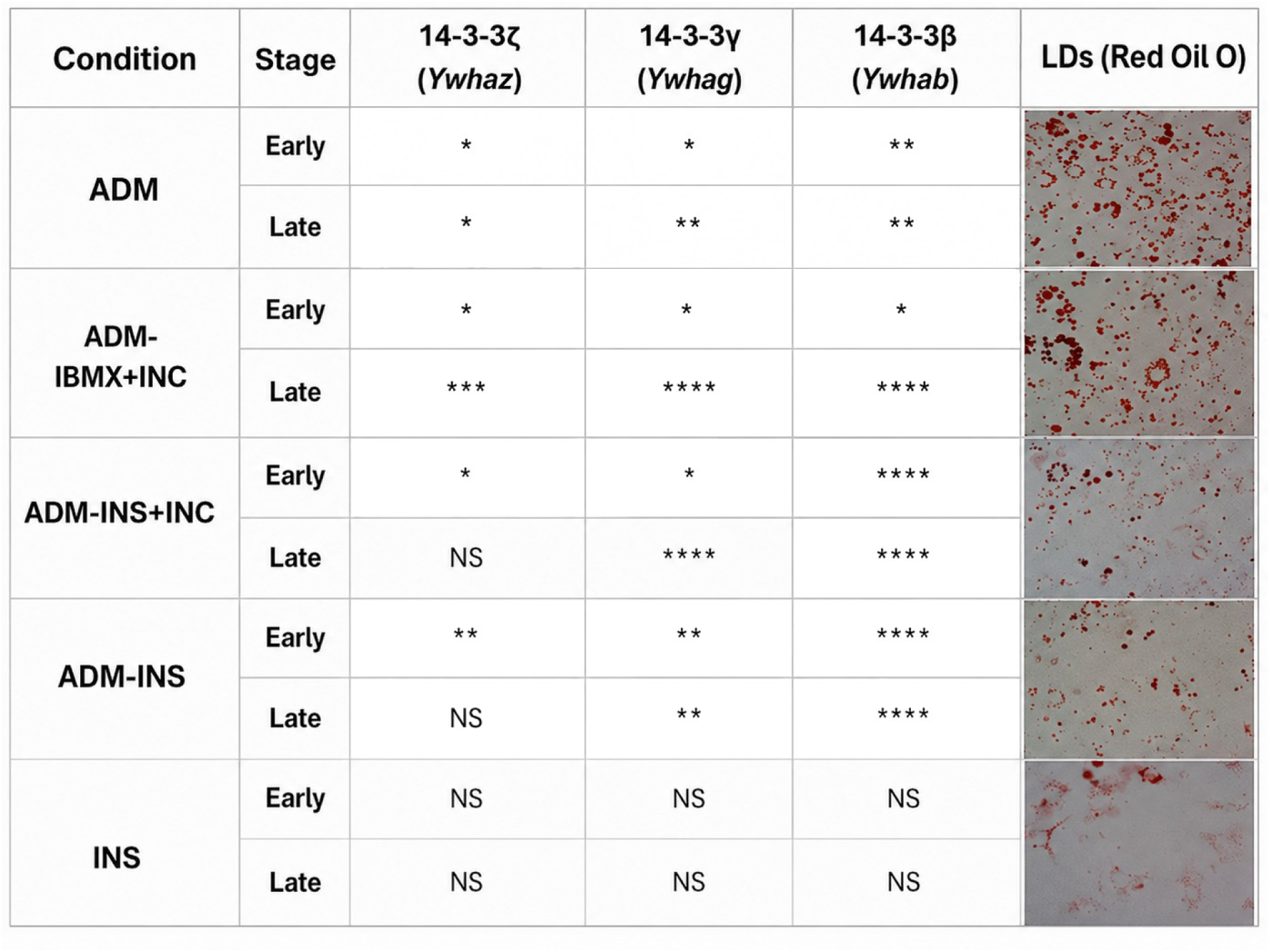
Effect of different differentiation media on lipid accumulation and 14-3-3 paralog expression during 3T3-L1 adipocyte differentiation. 3T3-L1 cells were differentiated using various adipogenic media containing combinations of dexamethasone, rosiglitazone, IBMX, GLP-1, and insulin, as indicated. Lipid accumulation was evaluated on day 7 by Oil Red O staining, showing higher lipid deposition under complete ADM and ADM supplemented with GLP-1 compared to other conditions. The table below shows the relative expression changes of 14-3-3 paralogs (γ, β and ζ) at early and late stages of differentiation, MDA in comparison to other conditions. NS: not significant.

This figure compares how different differentiation media influence lipid accumulation and 14-3-3 paralog expression during 3T3-L1 adipocyte differentiation. Oil Red O staining shows that full adipogenic medium (ADM) and ADM supplemented with incretin promote strong lipid accumulation, while insulin alone induces minimal differentiation. The accompanying statistical table indicates that 14-3-3*γ* and *β* are significantly upregulated at late differentiation stages under complete ADM conditions but remain unchanged with insulin alone. Overall, the results demonstrate that full hormonal induction is required for effective adipocyte differentiation and activation of specific 14-3-3 paralogs.

Overall, adipogenic medium (ADM) drives a clear late-stage upregulation of all 14-3-3 isoforms, accompanied by strong lipid droplet formation. This effect is further enhanced when IBMX plus incretin are included (ADM–IBMX+INC), which shows even stronger late-stage increases and more abundant lipid accumulation in the Oil Red O images. Conditions combining insulin with incretin (ADM–INS+INC) or insulin alone (ADM–INS) produce a more mixed response, with weaker or more variable induction of 14-3-3 expression and reduced lipid accumulation compared to ADM. In contrast, insulin alone (INS) shows no significant changes across all isoforms and no visible lipid droplet formation, indicating that insulin by itself is insufficient to trigger adipogenic differentiation in this system.

### 6. Silencing of 14-3-3β or 14-3-3γ differentially affects lipid droplet accumulation during adipogenic differentiation

To investigate the specific contribution of individual 14-3-3 paralogs during adipogenic differentiation, we used previously established cellular models described in Frontini et al. and Rivera et al. In these models, 14-3-3β (*Ywhab*) or 14-3-3γ (*Ywhag*) expression was selectively knockdown. Figure 7 shows confocal microscopy images of cells expressing shRNAs targeting *Ywhag* or *Ywhab* in mixed cultures with WT cells, together with the quantification of lipid droplet area per cell (μm²) in undifferentiated and adipogenically differentiated cells. We observed that lipid droplet accumulation increased after adipogenic differentiation in both knockdown conditions compared with untreated (UT) cells. However, comparison between differentiated WT cells and cells with *Ywhag* or *Ywhab* knockdown revealed different phenotypes. While *Ywhab* silencing resulted in reduced lipid accumulation compared with WT cells, *Ywhag* silencing produced the opposite effect, leading to increased lipid droplet accumulation.

**Figure 7.**
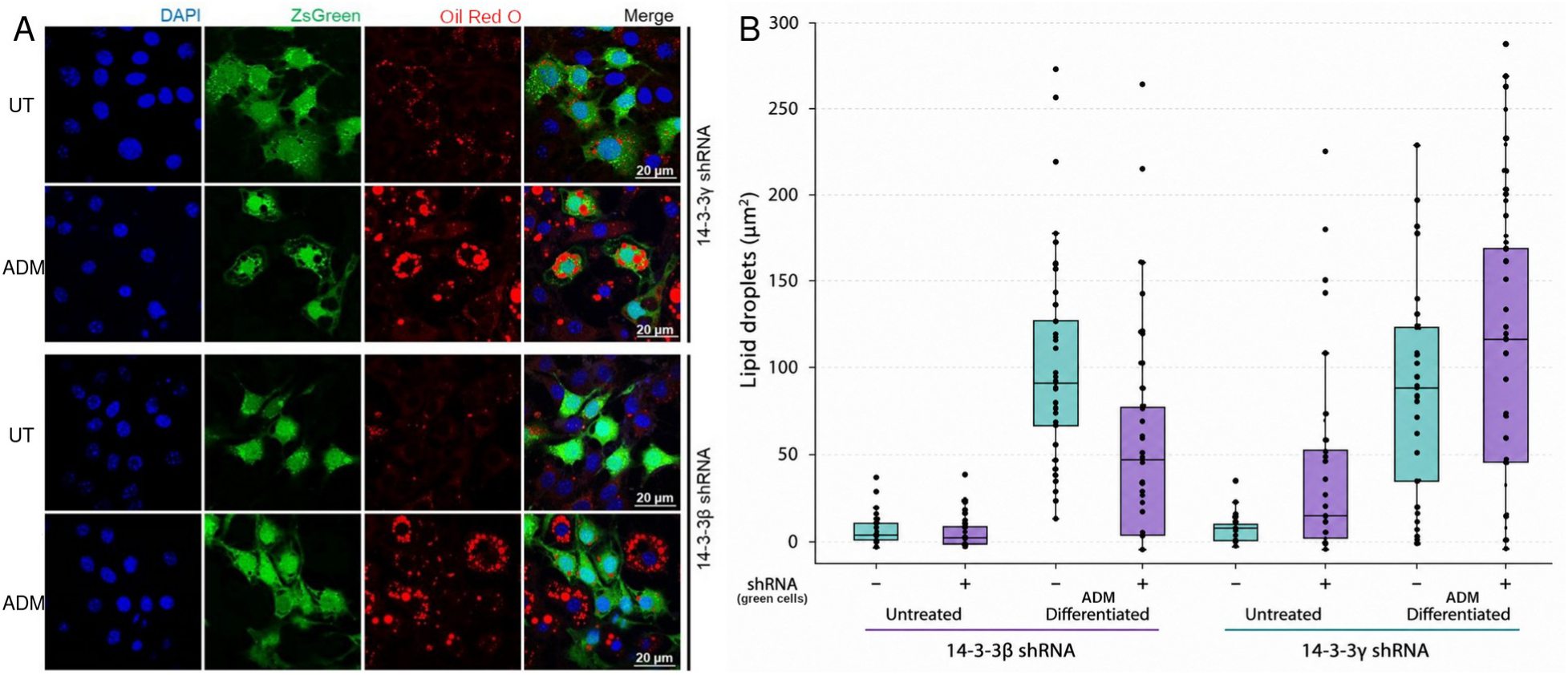
Knockdowm of Ywhag and Ywhab deferentially affects lipid droplet accumulation during adipogenic differentiation. (A) Confocal microscopy images of cells expressing shRNAs targeting Ywhag or Ywhab in a mixed culture with WT cells. Untreated (UT) conditions or after adipogenic differentiation (ADM). Nuclei were stained with DAPI (blue), shRNA-expressing cells were identified by ZsGreen fluorescence (green), and neutral lipid droplets were stained with Oil Red O (red). Merge images show the overlap of all channels. Scale bars, 20 μm. (B) Quantification of lipid droplet area per cell (μm²) in untreated and differentiated cells expressing control (−) or 14-3-3-targeting (+) shRNAs. Each dot represents an individual cell; boxes indicate the interquartile range (25th–75th percentiles), the center line represents the median, and whiskers extend to 1.5 × the interquartile range. Lipid droplet accumulation increased markedly following adipogenic differentiation in both knockdown conditions, with 14-3-3γ silencing producing the greatest increase in differentiated cells. Data are representative of at least three independent experiments.

## DISCUSSION

3T3-L1 preadipocytes can be efficiently induced to differentiate into adipocytes using a differentiation cocktail containing 0.5 mM IBMX, 0.25 µM dexamethasone, and 1 µg/mL insulin in DMEM supplemented with 10% FBS. Under these conditions, mature adipocytes typically develop within approximately two weeks. Differentiation protocols may also include 2 µM rosiglitazone, a thiazolidinedione antidiabetic drug that acts as an insulin sensitizer, with or without prolonged IBMX treatment, to enhance adipogenic differentiation (Zhao *et al*., 2019). In addition to these classical agents, alternative approaches to induce adipogenesis have been less explored. For instance, cAMP induction -typically achieved with IBMX- can also be triggered by incretins. These hormones are secreted by the gastrointestinal tract in response to nutrient ingestion, leading to enhanced glucose-stimulated insulin secretion (Baggio S Drucker, 2007). 3T3-L1 cells express functional incretin receptors, and it has been demonstrated that incretins are involved in lipid metabolism (Baggio S Drucker, 2007). Using a cAMP sensor plasmid, we show that 3T3-L1 cells respond to incretins (Fig. 3). Replacing IBMX with incretins resulted in adipocytes containing a greater number of smaller lipid droplets (Fig. 5). Although the cells responded to this treatment, no differences in 14-3-3 protein expression were observed between the two differentiation conditions.

The importance of the 14-3-3 protein family in adipogenesis is often underappreciated, and some paralogs have been considered housekeeping proteins due to their invariant levels during differentiation. However, in the present study, we identified novel regulatory roles for two of the seven 14-3-3 paralogs in murine preadipocyte 3T3-L1 cells, which complement those previously observed for the paralog *ζ* (Lim 2015, 2016). For the paralog 14-3-3*γ* , we observed a high dependence to insulin to increase its levels during differentiation, showing that insulin is required to enhance the expression of the mRNA of this paralog (Fig. 2B). Those conditions where insulin was absent there is not difference in the 14-3-3*γ* mRNA between day 3 and day 7 of differentiation. Western blot experiment using specific antibodies show that at protein level insulin has not modified its levels (Fig. 4A).

On the other hand, 14-3-3*β* responds differently to the addition of insulin in the differentiation medium. The absence of insulin significantly increases 14-3-3 *β* mRNA levels compared to conditions such as MDA or MDA-IBMX+incretins, where insulin is present (Fig 2C). This marked difference between insulin-free and insulin-containing conditions is also evident at the protein level, as Western blot experiments show a concentration-dependent effect of insulin on 14-3-3*β* expression (Fig. 4B).

The influence of insulin on the expression of 14-3-3 proteins has been previously reported (Lim *et al*., 2015; Chan *et al*., 2019). These studies were focused on 14-3-3*ζ*. However, the cell culture conditions used in Chan studies differed markedly from those applied in the present work. In the earlier studies, 3T3-L1 cells were cultured in low-glucose DMEM supplemented with calf serum, whereas in our experiments, high-glucose DMEM and fetal bovine serum were used.

By analyzing the degree of differentiation in Oil Red O–stained images and comparing it with 14-3-3 paralog expression (Figs. 5 and 6), we observed that the expression of 14-3-3*β*, which is inhibited by insulin, correlates with reduced differentiation, as earlier expression was associated with fewer lipid droplets in differentiated 3T3-L1 cells. Under insulin-free conditions, such as ADM–insulin+GLP-1 or ADM–insulin, we observed increased 14-3-3 *β* expression during both early and late adipogenesis. In the absence of insulin, Oil Red O– stained images showed fewer lipid droplets, correlating with the strong early expression of 14-3-3*β*.

Overall, our results show the expression of 14-3-3*γ*, *β* and *ζ* are coordinated during adipogenesis and appear to be important determinants in adipogenesis. Based on comparative analysis, we conclude that 14-3-3 have potentially novel physiological roles in the regulation of early and late adipogenesis and could be interesting therapeutic targets.

## Acknowledgements

SdV, SM and ADG were fellows of the National Research Council of Argentina (CONICET), JNA is a fellow of ANPCyT. SdV was also supported by an IUBMB Wood-Whelan Fellowship. MU and DMB are researchers of the same institution. This report was supported by the following grants: PIP-0118 (CONICET), PICT’21-0180 ANPCyT. GEL was supported by CIHR Project (PJT-153144) and NSERC Discovery (RPGIN-2017-05209) grants, as well as funding from the Cardiometabolic Health Diabetes, and Obesity (CMDO) Research Network (Jean- Davignon Young Investigator Award) and the Banting Research Foundation (Discovery Award). GEL holds the Canada Research Chair in Adipocyte Development.

